# Cannabinoid Biosynthesis using Noncanonical Cannabinoid Synthases

**DOI:** 10.1101/2020.01.29.926089

**Authors:** Maybelle K. Go, Kevin Jie Han Lim, Wen Shan Yew

## Abstract

We have found enzymes from the berberine-bridge enzyme (BBE) superfamily (IPR012951) that catalyze the oxidative cyclization of the monoterpene moiety in cannabigerolic acid (CBGA) to form cannabielsoic acid B (CBSA). The enzymes are from a variety of organisms and are previously uncharacterized. This is the first report that describes enzymes that did not originate from the *Cannabis* plant that catalyze the production of cannabinoids. Out of 72 homologues chosen from the enzyme superfamily, six orthologues were shown to accept CBGA as a substrate and catalyze the biosynthesis of CBSA. The six enzymes discovered in this study are the first report of heterologous expression of BBEs that did not originate from the *Cannabis* plant that catalyze the production of cannabinoids using CBGA as substrate. This study details a new avenue for discovering and producing natural and unnatural cannabinoids.

---

*Cannabis sativa* contains at least 113 cannabinoid compounds; these cannabinoids exhibit a range of unique pharmacological and therapeutic properties. Despite the report of biosynthesis of tetrahydrocannabinolic acid (THCA) and cannabidiolic acid (CBDA) using yeast ^1^, the biosynthetic pathways of a majority of these cannabinoids remain largely unknown, making it difficult for medicinal cannabinoids to be produced in a stable and sustainable manner. Chemical synthesis of these cannabinoids remains a challenge due to their complex chemical structures. Most of the well-known and studied cannabinoids are derived from cannabigerolic acid (CBGA). Six of the products, THCA, CBDA, tetrahydroisocannabinolic acid, cannabicyclolic acid, cannabichromenic acid, and cannabicitranic acid are structural isomers (C_22_H_30_O_4_); only cannabielsoic acid B (CBSA) is unique due to an additional oxygen atom (C_22_H_30_O_5_) (Extended Data Figure 1).

The biosynthesis of cannabinoids from CBGA is catalyzed by cannabinoid synthases. The enzymes are characterized as flavin adenine dinucleotide (FAD) – dependent berberine bridge enzymes that catalyze the oxidative cyclization of the monoterpene moiety in CBGA. The structure of THCA synthase (Uniprot ID Q8GTB6, PDB ID 3VTE) was elucidated by Kuroki and his associates in 2012 ^2^. Cannabidiolic acid synthase was functionally expressed in yeast by Stehle and his associates in 2017 ^3^. The known cannabinoid synthases share approximately 80% sequence identity.

THCA synthase is an enzyme of the berberine-bridge enzyme family (IPR012951). The family contains about 21,000 sequences, of which 37 are experimentally annotated and reviewed. Using the EFI-EST tool ^4^ to generate a sequence-similarity network (SSN) of this family, it is possible to segregate the proteins into iso-functional clusters. This will place uncharacterized enzymes in sequence-function context with proteins that have been previously characterized from reliable experiments ^5,6^. Using this information, it is possible to find potential enzymes from other organisms that will catalyze the oxidative cyclization of the monoterpene moiety in CBGA. In this study, 72 homologues were chosen from the generated SSN that may potentially accept CBGA as a substrate and catalyze a FAD-dependent oxidative cyclization reaction (Figure 1).

**Figure 1.**
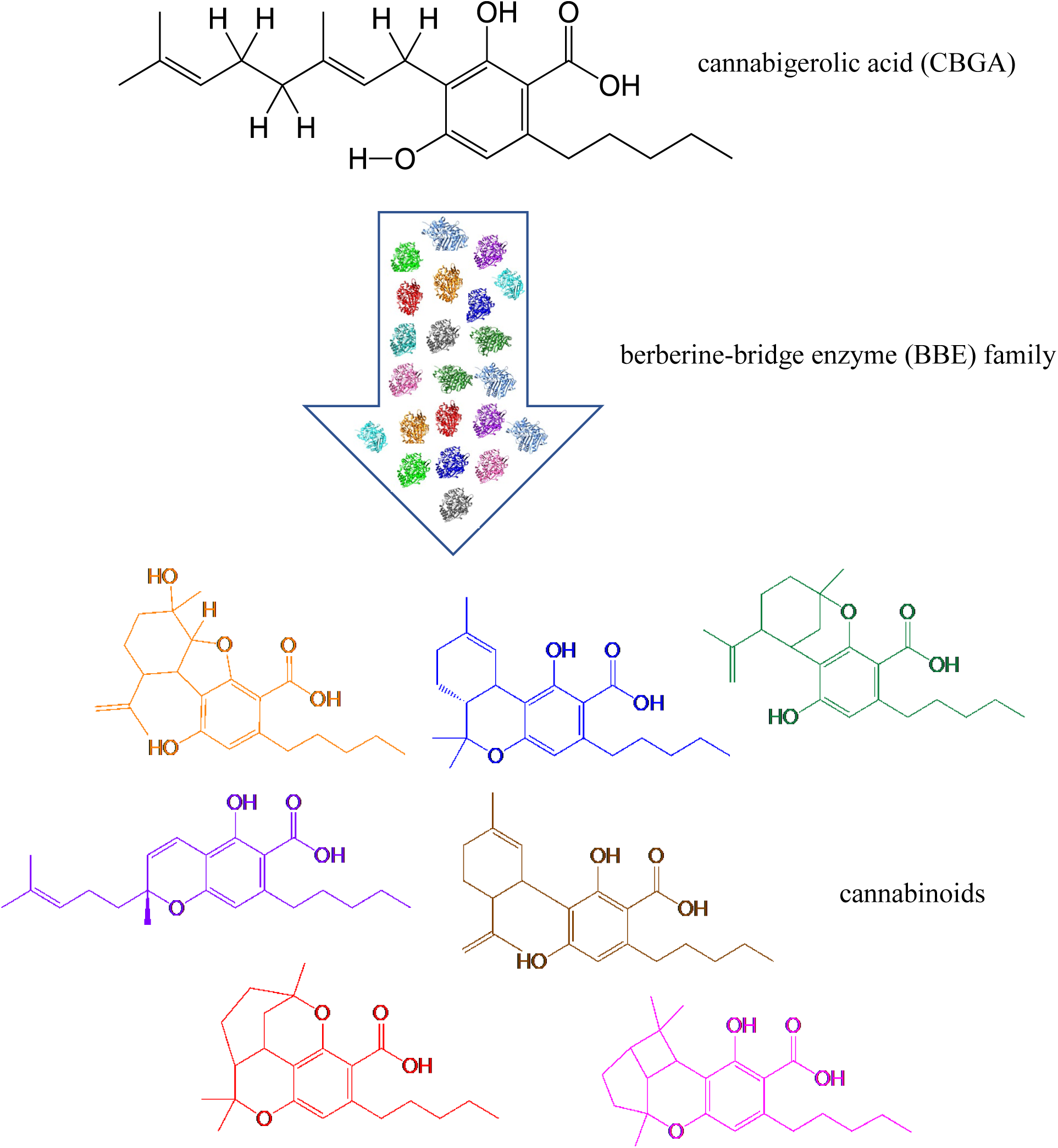
Production of cannabinoids catalyzed by noncanonical cannabinoid synthases from the berberine-bridge enzyme family.

The SSN generated approximately 4,500 nodes with 300,000 edges (Figure 2). A small number of homologues (72 homologues) were chosen based on either its similarity to THCA synthase or its difference. The goal is to create a small yet chemically diverse library to discover novel enzyme activity. The hypothesis of the study follows that enzymes similar to THCA synthase may catalyze the production of cannabinoids similar to THCA; enzymes “different” from THCA synthase may catalyze the production of novel cannabinoids.

**Figure 2.**
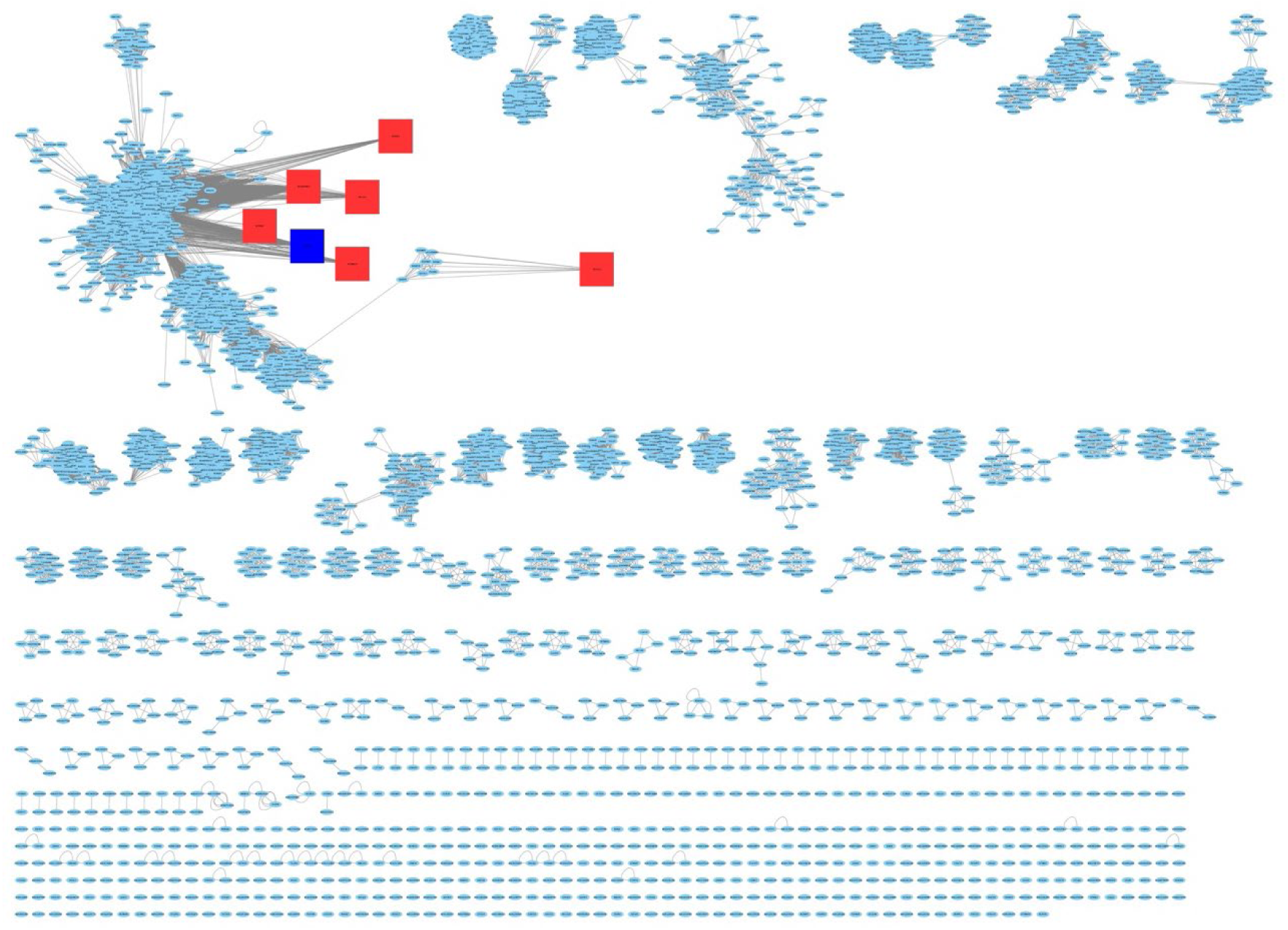
Full SSN of BBEs (IPR012951). The six orthologues are denoted by the red squares. THCA is denoted by the blue square. The six orthologues and THCA are not related (no edge connects them) but are part of a large BBE cluster. Image generated using Cytoscape ^7^.

This report provides new enzymes that can advance the molecular production of cannabinoid molecules that exist in Nature, but wherein there are no known molecular tools or enzymes to biosynthesize them. Conceptually, the work also advances the approach to biosynthesize natural products that otherwise are refractory to bioproduction due to the lack of suitably identified enzymes in the biosynthetic pathways.

Annotated enzymes from *Cannabis sativa* did not produce detectable amounts of a cannabinoid: it is possible that the signal peptide impeded the heterologous expression in *S. cerevisiae* ^3^. The majority of the homologues chosen (92% of sequences) did not produce any detectable amounts of a cannabinoid, which may indicate that the active sites of BBEs are not as promiscuous as other enzyme superfamilies. Since the organisms chosen for this study are all eukaryotes, it is also possible that the annotated sequences contain introns, which may render the synthesized genes inactive. Surprisingly, six orthologues were discovered to utilize CBGA to catalyze the production of CBSA. A close inspection of the SSN (Figure 2) indicates that the orthologues are not related to each other (no edge connects them), nor related to THCA synthase. Serendipitously, they are all part of the biggest BBE cluster. Table 1 shows the Uniprot IDs of orthologues and the organisms where they originated from.

**Table 1.** Berberine-bridge enzymes that catalyze the production of cannabinoids.

| Uniprot ID | Organism |
| --- | --- |
| D7MMG9 | *Arabidopsis lyrata subsp. lyrata* |
| M4DIE5 | *Brassica rapa subsp. pekinensis* |
| A0A1J6KPK0 | *Nicotiana attenuata* |
| M5X864 | *Prunus persica* |
| O64745 | *Arabidopsis thaliana* |
| P93479 | *Papaver somniferum* |

The enzymes D7MMG9, M4DIE5, A0A1J6KPK0, and M5X864 are uncharacterized proteins. The sequences are annotated as BBE-like enzymes based on sequence homology. O64745 from *Arabidopsis thaliana* was detected at the transcript level and is proposed to catalyze BBE activity in monolignol metabolism^8^. P93479 from *Papaver somniferum* (opium poppy) was also detected at the transcript level and is proposed to be involved in formation of *(S)*-scoulerine *via* the oxidative cyclization of the *N-*methyl moiety of *(S)*-reticuline.

The orthologues share about 35% sequence identity to THCA synthase (Extended Data Figure 2). The orthologues were submitted to the Protein Homology/analogY Recognition Engine V 2.0 (Phyre2) to generate homology models ^9^. The final model for each orthologue was used as the template to compare against THCA synthase. The two structures were aligned using the MatchMaker function in UCSF Chimera ^10^ to identify analogous positions in the orthologues that are critical for cannabinoid synthase activity (Extended Data Figure 3). On average the root-mean-square deviation (RMSD) score of the different alignments is less than 1 Å.

Five residues in THCA synthase are critical for its activity ^2^ – two catalytic residues, His-114 and Cys-176, which are covalently bonded to FAD, and His-292, Tyr-417, and Tyr-484. Table 2 shows critical residues in THCA synthase and the analogous residues in the orthologues (Extended Data Figure 4). Four of the orthologues – A0A1J6KP0, M5X864, O64745, and P93479, contain both catalytic residues and also the corresponding Tyr-484 residue, whilst His-292 and Tyr-417 were not conserved. Amongst these four orthologues, M5×864 retained amino acid chemical functionality, *vis-a-vis* His-292 altered to Gln-241 and Tyr-417 changed to Thr-378. The other three orthologues did not retain amino acid chemical functionality: His-292 was altered to a hydrophobic residue (Val-291 or Leu-286) or an acidic residue (Glu-294) and Tyr-417 was changed to an Asn residue (Asn-408, Asn-414 and Asn-394, respectively). Surprisingly, D7MMG9 and M4DIE5 lacked either one or both catalytic residues. The other critical residues were also not conserved. Nevertheless, these two orthologues have detectable cannabinoid synthase active to catalyze the production of cannabielsoic acid (CBSA).

**Table 2.** Critical residues in THCA synthase with the corresponding analogous residues in the orthologues. The proteins are identified by their UNIPROT ID. Residues are labelled using the single letter amino acid code.

|  | UNIPROT ID |  |  |  |  |  |  |
| --- | --- | --- | --- | --- | --- | --- | --- |
|  | 3VTE | D7MMG9 | M4DIE5 | A0A1J6KP0 | M5X864 | O64745 | P93479 |
| Residue | H114 | H73 | - | H112 | H54 | H116 | H108 |
|  | C176 | - | - | C176 | C116 | C178 | C170 |
|  | H292 | T232 | L98 | V291 | Q241 | E294 | L286 |
|  | Y417 | T345 | N211 | N408 | T378 | N414 | N394 |
|  | Y484 | F417 | Y278 | Y477 | Y445 | Y483 | H463 |

The six enzymes discovered in this study are the first report of heterologous expression of BBEs that did not originate from the *Cannabis* plant that catalyze the production of cannabinoids using CBGA as substrate. It is recognized that the study only explored a very limited portion of this enzyme family (72 out of 21,000; 0.34%); the corollary expectation follows that other enzymes within this family will accept either CBGA or/as well as the shorter analogs (with *n-*butyl, *n*-propyl, ethyl and methyl side chains) as substrates. This study delineates a new avenue for the discovery and the biosynthesis of natural and unnatural cannabinoids.

The structure of cannabielsoic acid is shown in Figure 3; HR-MS (ESI, negative mode): [M-H]^-^, m/z = 373.2008 (experimental); m/z = 373.2020 (theoretical). The fragments, [M–CO_2_]^-^, [M– CH_3_]^-^, and [M–C_3_H_5_]^-^, were also detected with m/z = 329, 358, and 332 respectively. The assignments of ^1^H- and ^13^C-NMR spectra are shown in Table 3 and the Heteronuclear Multiple Quantum Coherence (HMQC) spectra in Extended Data Figure 5. The chemical shifts were assigned based on two previous works. In 1974, Shani and Mechoulam isolated CBSA from hashish and elucidated the structure from spectral data ^11^. Unfortunately, only proton chemical shifts were assigned. In 2004, work by Verpoorte and his associates determined NMR assignments (^1^H and ^13^C) of major cannabinoids isolated from the plant ^12^. Based on the published data of major cannabinoids such as cannabidiolic acid, ^1^H- and ^13^C-NMR assignments for CBSA were determined.

**Table 3.**
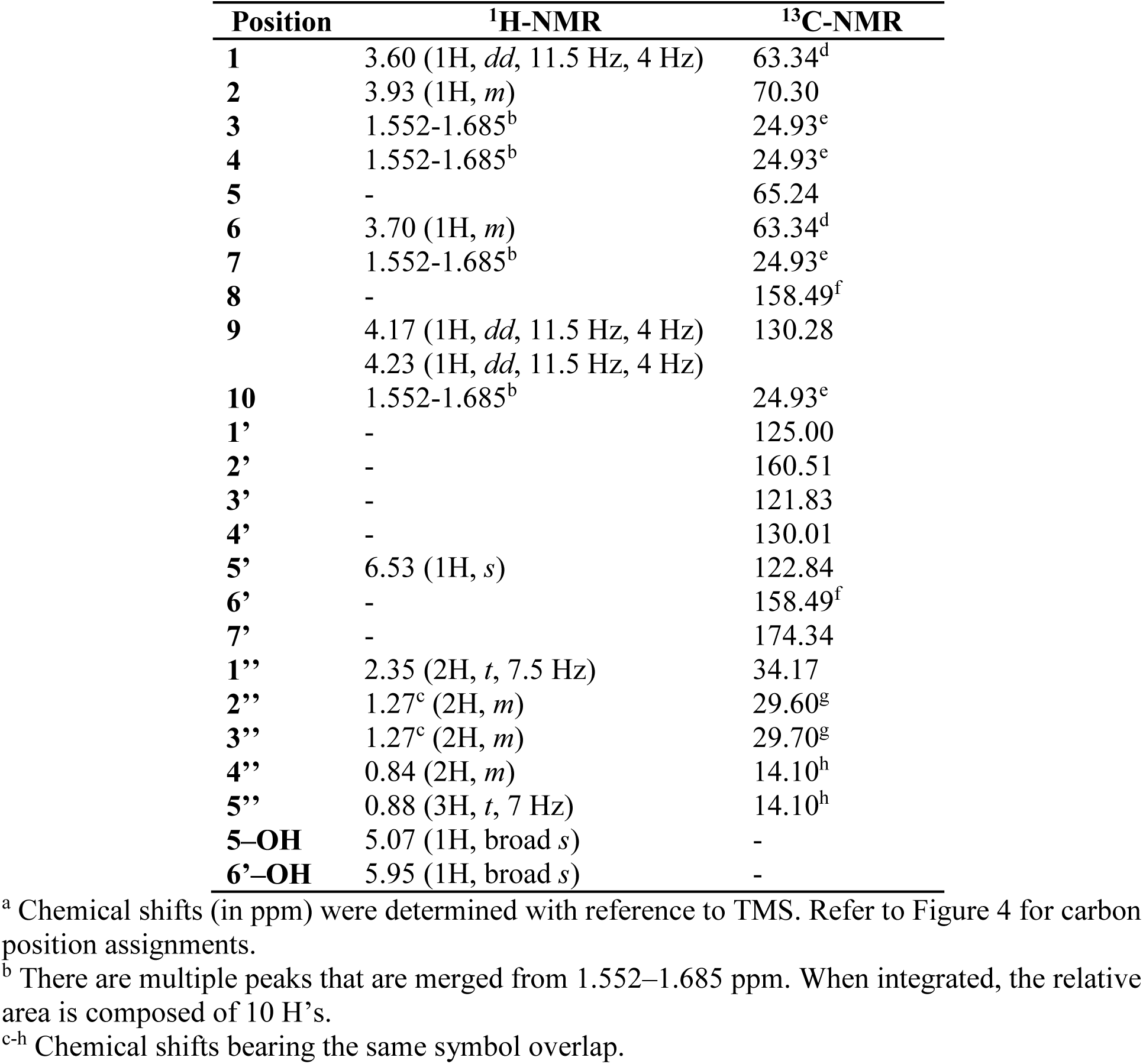
^1^H- and ^13^C-NMR assignments for CBSA in CDCl_3_ ^a^.

| Position | <sup>1</sup>H-NMR | <sup>13</sup>C-NMR |
| --- | --- | --- |
| **1** | 3.60 (1H, *dd*, 11.5 Hz, 4 Hz) | 63.34<sup>d</sup> |
| **2** | 3.93 (1H, *m*) | 70.30 |
| **3** | 1.552-1.685<sup>b</sup> | 24.93<sup>e</sup> |
| **4** | 1.552-1.685<sup>b</sup> | 24.93<sup>e</sup> |
| **5** | - | 65.24 |
| **6** | 3.70 (1H, *m*) | 63.34<sup>d</sup> |
| **7** | 1.552-1.685<sup>b</sup> | 24.93<sup>e</sup> |
| **8** | - | 158.49<sup>f</sup> |
| **9** | 4.17 (1H, *dd*, 11.5 Hz, 4 Hz) | 130.28 |
|  | 4.23 (1H, *dd*, 11.5 Hz, 4 Hz) |  |
| **10** | 1.552-1.685<sup>b</sup> | 24.93<sup>e</sup> |
| **1'** | - | 125.00 |
| **2'** | - | 160.51 |
| **3'** | - | 121.83 |
| **4'** | - | 130.01 |
| **5'** | 6.53 (1H, *s*) | 122.84 |
| **6'** | - | 158.49<sup>f</sup> |
| **7'** | - | 174.34 |
| **1''** | 2.35 (2H, *t*, 7.5 Hz) | 34.17 |
| **2''** | 1.27<sup>c</sup> (2H, *m*) | 29.60<sup>g</sup> |
| **3''** | 1.27<sup>c</sup> (2H, *m*) | 29.70<sup>g</sup> |
| **4''** | 0.84 (2H, *m*) | 14.10<sup>h</sup> |
| **5''** | 0.88 (3H, *t*, 7 Hz) | 14.10<sup>h</sup> |
| **5-OH** | 5.07 (1H, broad *s*) | - |
| **6'-OH** | 5.95 (1H, broad *s*) | - |

<sup>a</sup> Chemical shifts (in ppm) were determined with reference to TMS. Refer to Figure 4 for carbon position assignments.
<sup>b</sup> There are multiple peaks that are merged from 1.552–1.685 ppm. When integrated, the relative area is composed of 10 H's.
<sup>c-h</sup> Chemical shifts bearing the same symbol overlap.

**Figure 3.**
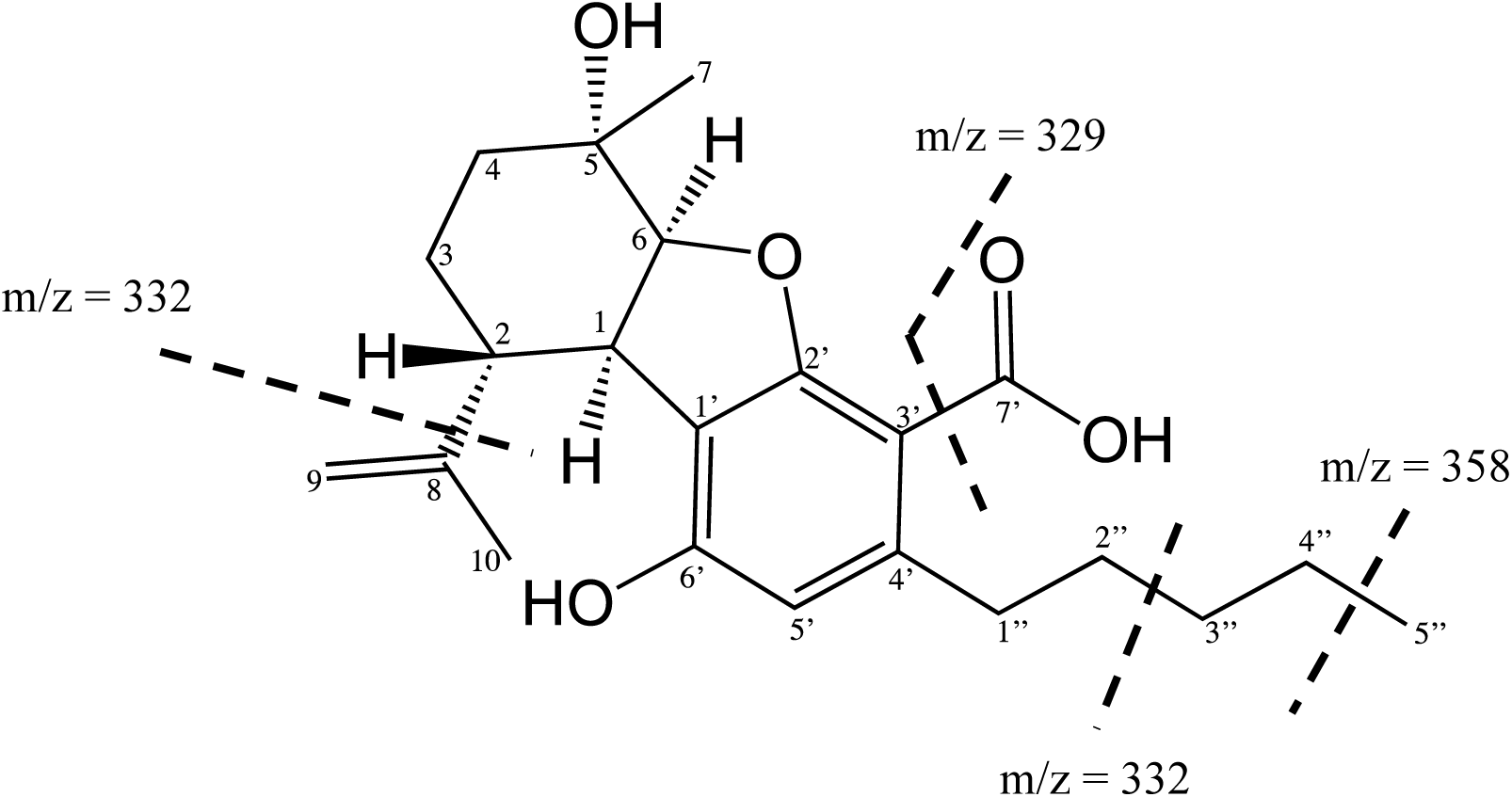
Chemical structure of cannabielsoic acid B (CBSA).

CBSA is different from the rest of the products due to an additional oxygen atom. It was previously reported to be an oxidative product from CBDA ^11^. Analogous to previously determined oxidative cyclization mechanisms of CBGA, we propose that it begins with the formation of the carbocation in the monoterpene moiety and the corresponding reduction of FAD to FADH_2_. The secondary carbocation formed rearranges to a more stable tertiary carbocation. Then, a proton is extracted from a terminal methyl group of the octadienyl chain by a general base, and a cascade of electron pair movements forms the cyclohexyl ring. Thereafter, we propose a nucleophilic attack of the carbocation intermediate (presumably by a nucleophile derived from a H_2_O molecule) to form 2,4-dihydroxy-3-[2-hydroxy-2-methyl-5-(2-propenyl)-cyclohexyl]-6-pentyl-benzoic acid. A second carbocation forms with a corresponding reduction of a second FAD molecule to FADH_2_ and a final cyclization step produces CBSA (Extended Data Figure 6).

We have discovered that enzymes from the berberine-bridge enzyme family (IPR012951) can catalyze the oxidative cyclization of the monoterpene moiety in CBGA to form CBSA. This is the first report of enzymes that did not originate from the *Cannabis* plant that catalyze the production of cannabinoids. This study demonstrated the attractive potential of curating sequence space, using tools such as SSNs, to discover uncharacterized enzymes and curate them based on sequence and structure. Further exploration may include expanding the screen towards the rest of the ∼21,000 sequences in the BBE family, as well as using the lesser known analogs of CBGA as substrates. We have also identified non-*Cannabis* enzymes within the BBE family sequences that are able to catalyze the oxidative cyclization of CBGA to form the structural isomers (C_22_H_30_O_4_) of THCA and CBDA; efforts are ongoing to characterize these THCA synthase and CBDA synthase orthologues. We believe that this discovery will aid in the production of cannabinoids in a stable and sustainable manner. Current work is focused on the characterization of these newly discovered cannabinoid synthase orthologues.

## Methodology

The sequence of THCA synthase was used as a query to determine the superfamily the enzyme belongs to (http://www.ebi.ac.uk/interpro/) ^13^. Thereafter, the EFI-EST tool (https://efi.igb.illinois.edu/efi-est/) was used to generate the SSN. A small number of homologues (72 homologues) were chosen to test if CBGA can be used as a substrate for these enzymes to catalyze the biosynthesis of a cannabinoid. The homologues were chosen using the following criteria: 1) Homologues found in *Cannabis sativa* and its related organisms such as *Nicotiana sp*.; 2) Homologues that are experimentally shown to exist either at the protein or transcript level; 3) Homologues from other plants that share less than 40% sequence identity to THCA synthase.

CBGA was purchased from Cayman Chemicals. All other chemicals used are of the highest purity that is required for the different experiments. The 72 homologues chosen from the SSN were codon optimized for yeast expression, synthesized, and cloned into pYES2 vector (Thermofisher).

The cloned genes were transformed into *Saccharomyces cerevisiae* BY4741 ^14,15^ by chemical transformation ^16^. The transformed cells were plated onto SC-URA-glucose plates and incubated for two days at 30 °C. Three single colonies were picked and grown in SC-URA-glucose media for 24 h. The cells were harvested and resuspended in SC-URA-galactose media and incubated for another 24 h. Thereafter, cells were harvested and resuspended in 100 mM citrate, pH 5.5 and 1 mM MgCl_2_. The cell wall was digested by addition of lyticase (0.5 ug) for 1 h at 37 °C. Glass beads (425-600 μm) were added and the cells were broken using a tissue homogenizer. The cell supernatant was clarified by centrifuge and used for the biosynthetic activity screen.

A 50-uL reaction mixture was prepared for the cannabinoid biosynthesis activity screen. The mixture contains 0.4 mM CBGA, 1.5 mM FAD, and 48 uL cell lysate. The reaction was incubated for 24 h under ambient conditions. The compounds were extracted using ethyl acetate, dried, and re-dissolved in acetonitrile. The presence of the cannabinoids was analyzed using the Agilent RapidFire 365 High-Throughput mass spectrometry system coupled with the Agilent 6495 LC-TQ. As mentioned previously, the potential cannabinoid products using CBGA have two different molecular formula - C_22_H_30_O_4_ and C_22_H_30_O_5_ with m/z values (ESI, negative mode) of 357.2 and 373.2 respectively. Corresponding control experiments in the absence of CBGA were also prepared.

Homologues that were determined to produce the cannabinoid compounds were re-analyzed using the Agilent 1290 Infinity HPLC coupled with the Agilent 6550 iFunnel Q-TOF high resolution mass spectrometer. This analysis will separate the different compounds present in the reaction as well as determine the exact mass of the compounds produced.

The yeast cells expressing homologues that were determined to produce cannabinoids by LC-MS analysis were grown on a larger scale to produce enough compounds for ^1^H-nuclear magnetic resonance (NMR) and ^13^C-NMR analysis. The steps performed are analogous to the smaller scale experiment mentioned previously. Samples were dissolved in CDCl_3_ and ^1^H-NMR and ^13^C - NMR spectra were recorded using a Bruker AVANCE 500 MHz NMR spectrometer at the Department of Chemistry, National University of Singapore.

## Acknowledgments

We thank Dr. Wei Zhe Teo for his assistance in using the Agilent RapidFire 365 High-Throughput mass spectrometry system coupled with the Agilent 6495 LC-TQ. We thank Ms. Yanhui Han for the NMR analytical service at the Department of Chemistry, National University of Singapore. This work was supported by the Synthetic Biology R&D Programme, National Research Foundation, Singapore.

## Author Contributions

M.K.G. and W.S.Y. conceived the study. M.K.G. and K.J.H.L. designed the biosynthetic screen platform and performed microbiological manipulations and extractions. M.K.G. designed and constructed the plasmids, performed the biosynthetic screening, mass spectrometry, and large-scale production of CBSA for NMR analysis. M.K.G. and W.S.Y. wrote the manuscript.

The authors declare no competing interests.

Reprints and permissions information is available at www.nature.com/reprints.

## Extended figures and tables

**Extended Data Figure 1.**
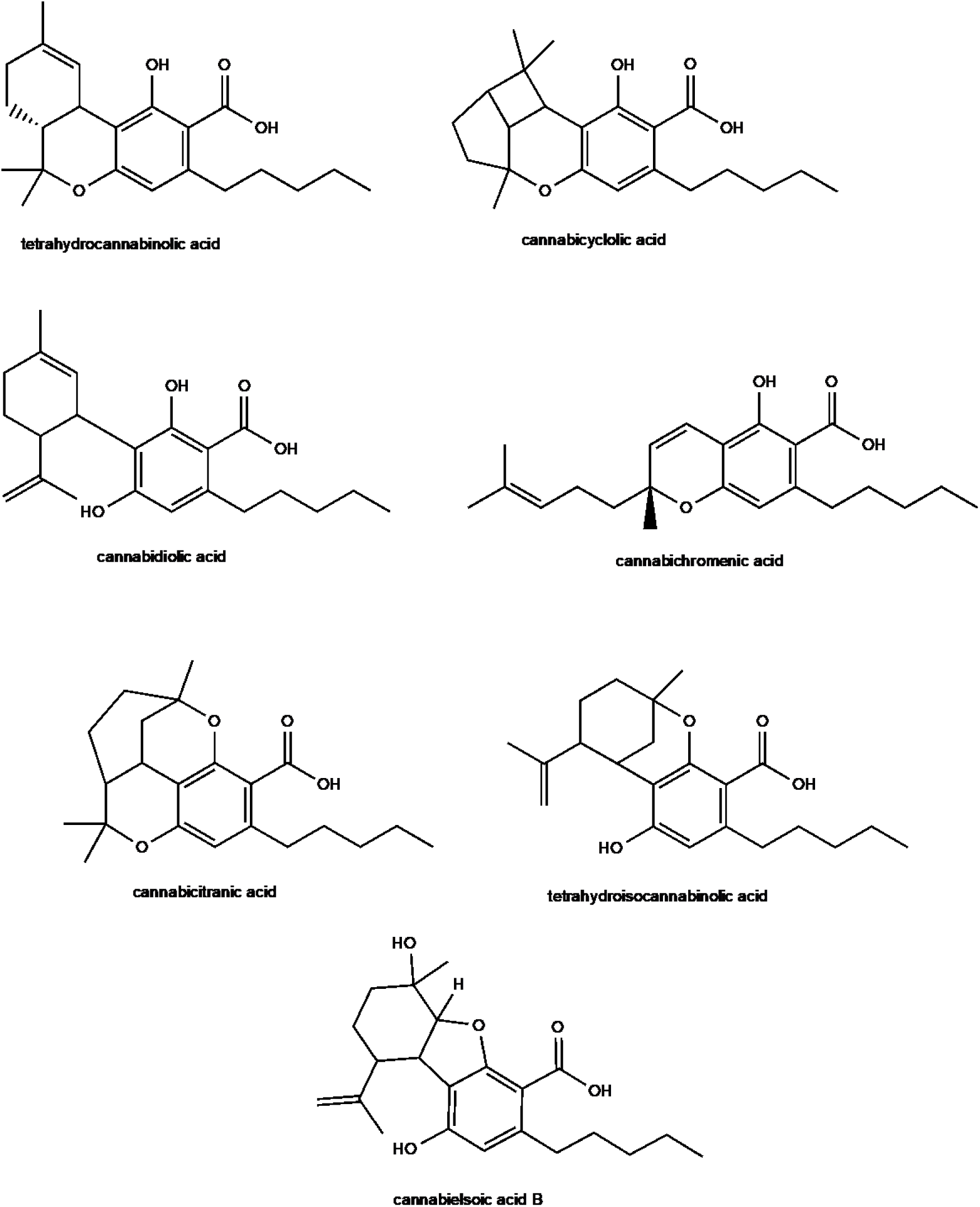
Structure of cannabinoids derived from CBGA.

**Extended Data Figure 2.**
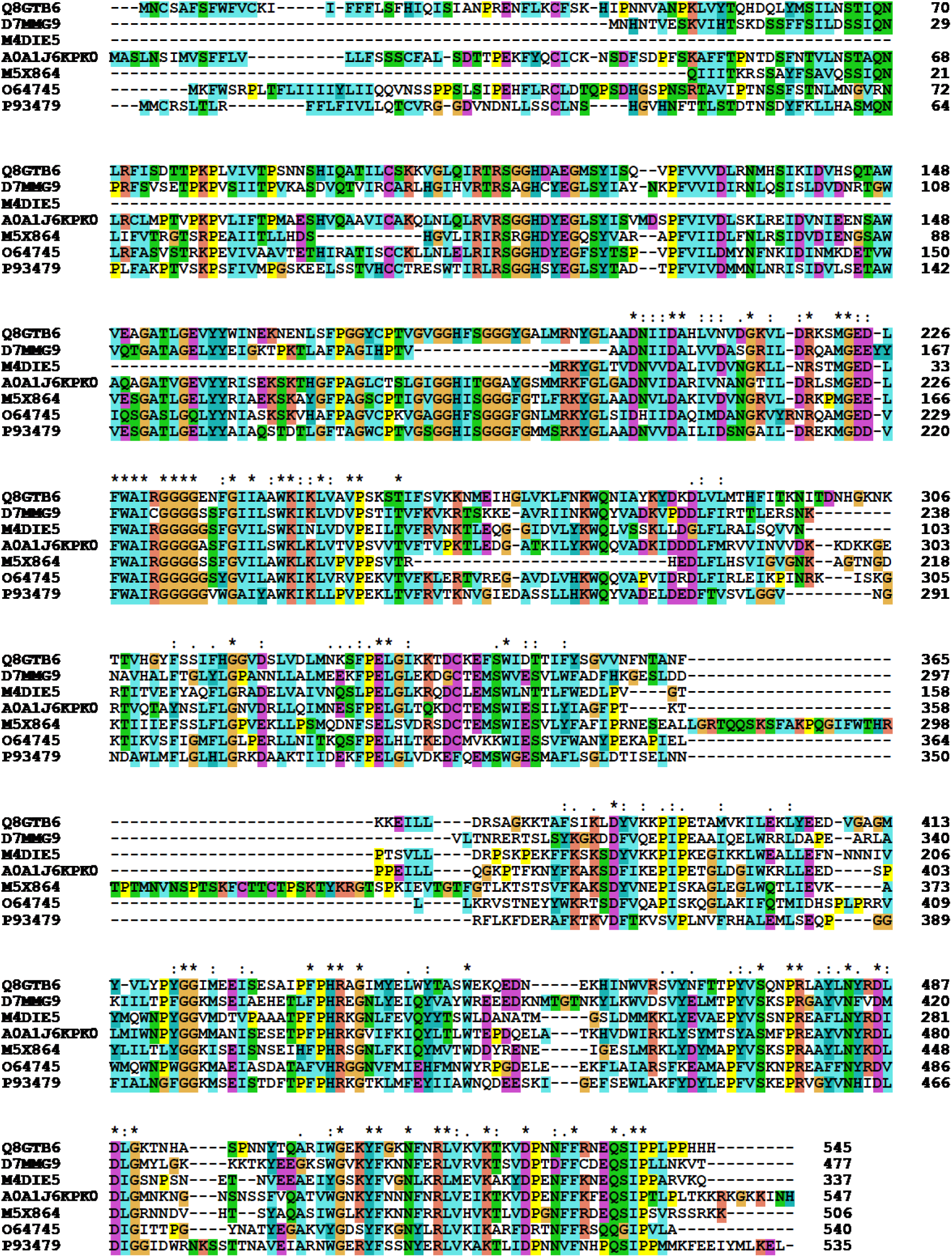
Sequence alignment of the six orthologues when compared to THCA synthase (Q8GTB6). Alignment generated using Clustal X ^16^.

**Extended Data Figure 3.**
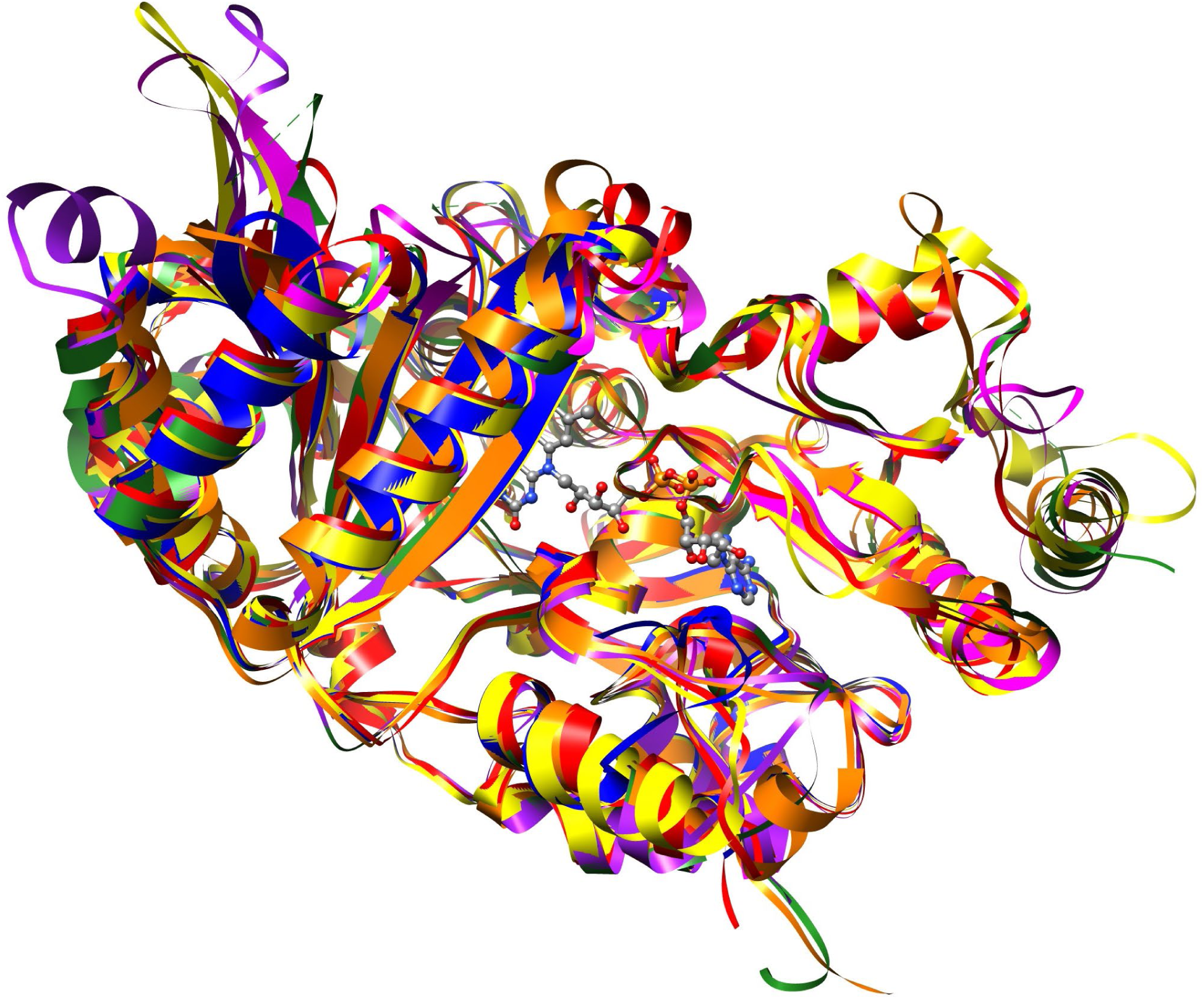
Ribbon model of 3VTE (green) aligned against all six orthologues -D7MMG9 (red), M4DIE5 (blue), A0A1J6KP0 (yellow), M5×864 (purple), O64745 (magenta), and P93479 (orange). The RMSD score is 0.986 Å. The models were generated using Phyre2 ^9^. The image was generated using UCSF Chimera 1.14 ^10^.

**Extended Data Figure 4.**
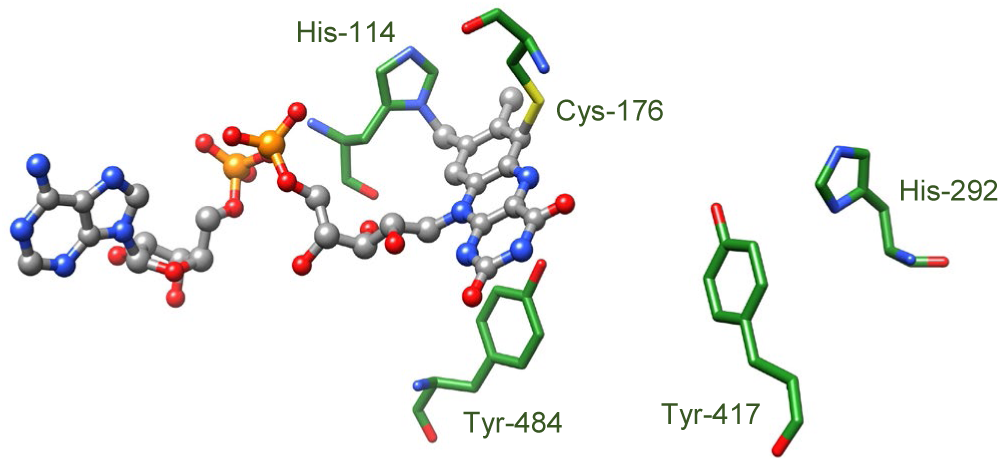
THCA synthase (PDB ID 3VTE) active site. The image was generated using UCSF Chimera 1.14 ^17^.

**Extended Data Figure 5.**
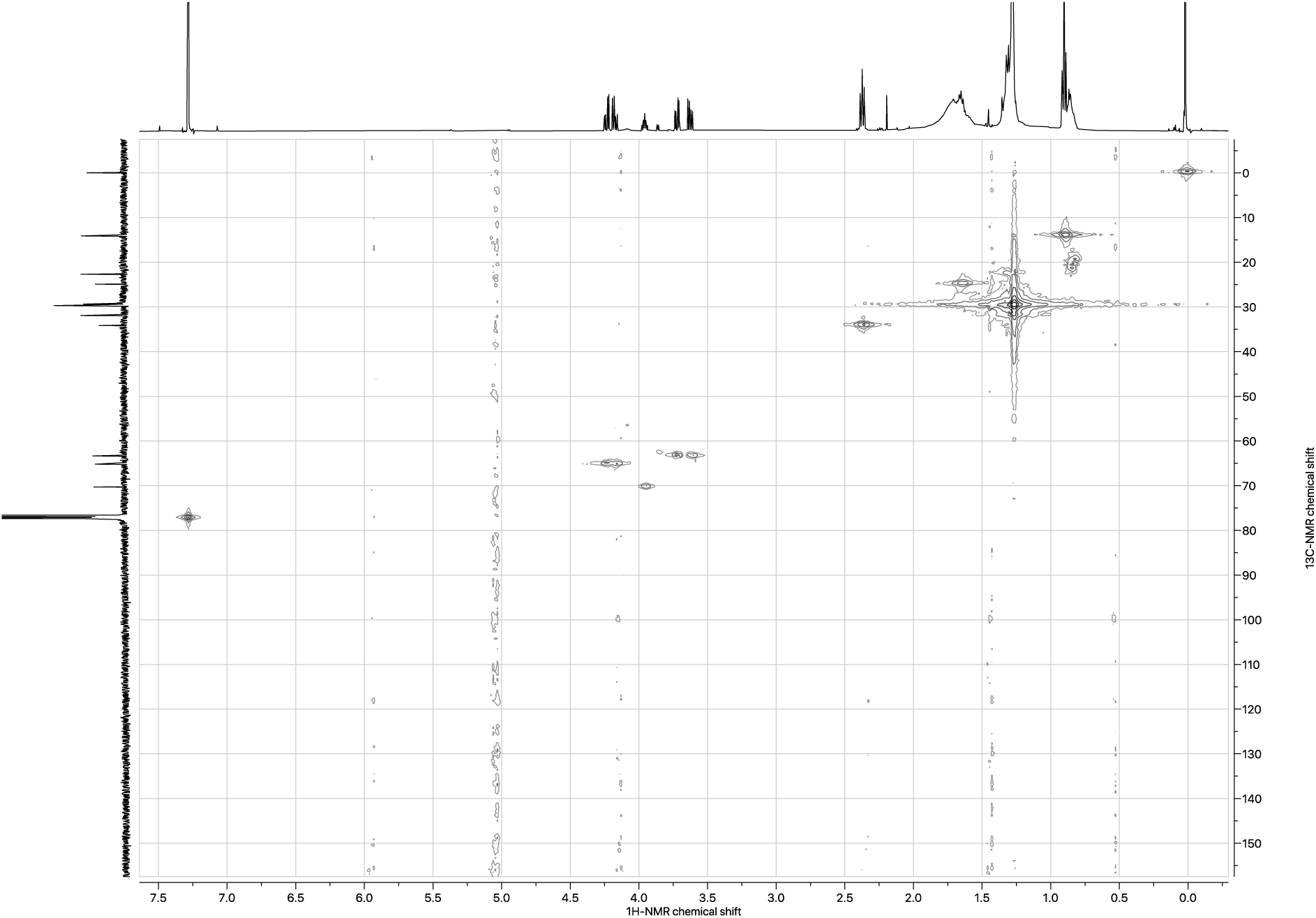
HMQC spectra of CBSA.

**Extended Data Figure 6.**
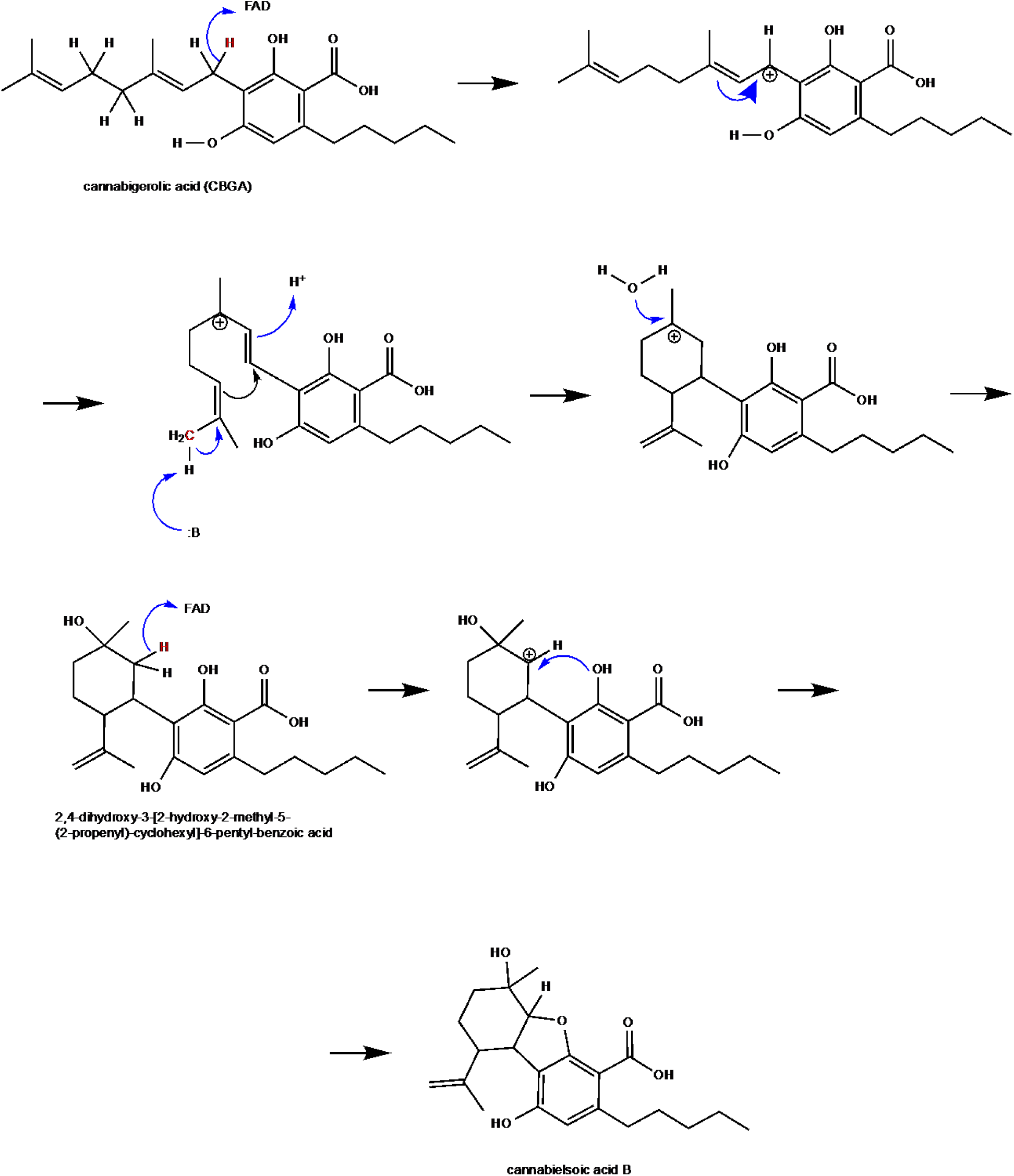
Proposed oxidative cyclization mechanism of CBSA formation from CBGA.

## References

1. Luo, X. et al. Complete biosynthesis of cannabinoids and their unnatural analogues in yeast. Nature 567, 123–126 (2019).

2. Shoyama, Y. et al. Structure and function of 1-tetrahydrocannabinolic acid (THCA) synthase, the enzyme controlling the psychoactivity of Cannabis sativa. Journal of molecular biology 423, 96–105 (2012).

3. Zirpel, B., Degenhardt, F., Martin, C., Kayser, O. & Stehle, F. Engineering yeasts as platform organisms for cannabinoid biosynthesis. Journal of biotechnology 259, 204–212 (2017).

4. Gerlt, J. A. et al. Enzyme Function Initiative-Enzyme Similarity Tool (EFI-EST): A web tool for generating protein sequence similarity networks. Biochim Biophys Acta 1854, 1019–1037 (2015).

5. Gerlt, J. A. Genomic Enzymology: Web Tools for Leveraging Protein Family Sequence-Function Space and Genome Context to Discover Novel Functions. Biochemistry 56, 4293–4308 (2017).

6. Zallot, R., Oberg, N. & Gerlt, J. A. The EFI Web Resource for Genomic Enzymology Tools: Leveraging Protein, Genome, and Metagenome Databases to Discover Novel Enzymes and Metabolic Pathways. Biochemistry (2019).

7. Shannon, P. et al. Cytoscape: a software environment for integrated models of biomolecular interaction networks. Genome research 13, 2498–2504 (2003).

8. Daniel, B. et al. Oxidation of Monolignols by Members of the Berberine Bridge Enzyme Family Suggests a Role in Plant Cell Wall Metabolism. The Journal of biological chemistry 290, 18770–18781 (2015).

9. Kelley, L. A., Mezulis, S., Yates, C. M., Wass, M. N. & Sternberg, M. J. E. The Phyre2 web portal for protein modeling, prediction and analysis. Nature Protocols 10, 845–858 (2015).

10. Pettersen, E. F. et al. UCSF Chimera--a visualization system for exploratory research and analysis. Journal of computational chemistry 25, 1605–1612 (2004).

11. Shani, A. & Mechoulam, R. Cannabielsoic acids: Isolation and synthesis by a novel oxidative cyclization. Tetrahedron 30, 2437–2446 (1974).

12. Choi, Y. H. et al. NMR assignments of the major cannabinoids and cannabiflavonoids isolated from flowers of Cannabis sativa. Phytochemical analysis : PCA 15, 345–354 (2004).

13. Mitchell, A. L. et al. InterPro in 2019: improving coverage, classification and access to protein sequence annotations. Nucleic acids research 47, D351–d360 (2019).

14. Winston, F., Dollard, C. & Ricupero-Hovasse, S. L. Construction of a set of convenient Saccharomyces cerevisiae strains that are isogenic to S288C. Yeast (Chichester, England) 11, 53–55 (1995).

15. Brachmann, C. B. et al. Designer deletion strains derived from Saccharomyces cerevisiae S288C: a useful set of strains and plasmids for PCR-mediated gene disruption and other applications. Yeast (Chichester, England) 14, 115–132 (1998).

16. Bergkessel, M. & Guthrie, C. in Methods in Enzymology Vol. 529 (ed Jon Lorsch) 311–320 (Academic Press, 2013).

17. Larkin, M. A. et al. Clustal W and Clustal X version 2.0. Bioinformatics (Oxford, England) 23, 2947–2948 (2007).

